# Thiamine pyrophosphate riboswitches in *Bacteroides* species regulate transcription or translation of thiamine transport and biosynthesis genes

**DOI:** 10.1101/867226

**Authors:** Zachary A. Costliow, Patrick H. Degnan, Carin K. Vanderpool

## Abstract

Thiamine (vitamin B_1_) and its phosphorylated precursors are necessary for decarboxylation reactions required in carbohydrate and branched chain amino acid metabolism. Due to its critical roles in central metabolism, thiamine is essential for human and animal hosts and their resident gut microbes. However, little is known about how thiamine availability shapes the composition of gut microbial communities and the physiology of individual species within those communities. Our previous work has implicated both thiamine biosynthesis and transport activities in the fitness of *Bacteroides* species. To better understand thiamine-dependent gene regulation in *Bacteroides*, we examined thiamine biosynthesis and transport genes in three representative species: *Bacteroides thetaiotaomicron, Bacteroides uniformis*, and *Bacteroides vulgatus*. All three species possess thiamine biosynthetic operons controlled by highly conserved *cis*-acting thiamine pyrophosphate (TPP) riboswitches. *B. thetaiotaomicron* and *B. uniformis* have additional TPP riboswitch-controlled operons encoding thiamine transport functions. Transcriptome analyses showed that each *Bacteroides* species had a distinct transcriptional response to exogenous thiamine. Analysis of transcript levels and translational fusions demonstrated that in *B. thetaiotaomicron*, the TPP riboswitch upstream of biosynthesis genes acts at the level of transcription, while TPP riboswitches upstream of transport operons work at the level of translation. In *B. uniformis* and *B. vulgatus*, TPP riboswitches work at the transcriptional level to control downstream operons. The varying responses to exogenous thiamine and use of varied regulatory mechanisms may play an important role in niche establishment by the Bacteroidetes in the complex and constantly shifting gut environment.

**Importance:** *Bacteroides* species are important and abundant members of human gut microbiome communities. Their activities in the gut are influenced by constant changes in nutrient availability. In this study, we investigated the genetic basis of thiamine (Vitamin B_1_) uptake and biosynthesis in three representative *Bacteroides* species. We found species-specific differences in the response to exogenous thiamine, and distinct mechanisms for regulation of uptake and biosynthesis gene expression. Our work implies that gut *Bacteroides* have evolved distinct strategies for making or acquiring an essential nutrient. These mechanisms may play an important role in the success of *Bacteroides* in establishing a niche within complex gut microbiome communities.

## Introduction

Thiamine pyrophosphate (TPP) is the active cofactor form of thiamine (Vitamin B_1_). Thiamine is required by all living organisms to perform a variety of metabolic activities. Thiamine-requiring pathways include central metabolism, branched chain amino acid biosynthesis, and nucleotide synthesis (1-4). Humans primarily acquire thiamine from their diet and convert it to the cofactor form, TPP (5, 6). In addition to dietary thiamine, there is some evidence that microbial communities in the gastrointestinal tract can provide thiamine to the host (7, 8). This thiamine may be supplied by resident gut microbes that possess the ability to synthesize thiamine and TPP *de novo* (9,10). A recent study suggested that certain pathogenic microbes can inhibit thiamine uptake by host intestinal epithelial cells (11). In addition, thiamine availability has been correlated with differences in composition of gut microbial communities (12). Thiamine excess and thiamine deficiency of the human host have been linked to different diseases, including Crohn’s Disease, Irritable Bowel Disease, and Dementia, which are also associated with altered gut microbiome structure and function (6,13,14). While these studies reveal interesting correlations that suggest a role for thiamine at the host-microbe interface, very little is known about how dominant members of the gut microbiota sense and respond to thiamine availability.

Work to characterize and understand thiamine biosynthesis and transport has primarily been done in model organisms including *Escherichia coli, Salmonella enterica*, and *Bacillus subtilis* (10, 15-18). Thiamine biosynthesis is carried out by a complex suite of enzymes through a conserved bifurcated pathway. The first key intermediates in TPP synthesis are 4-amino-2-methyl-5-diphosphomethylpyrimidine (HMP-PP) and 2-(2-carboxy-4-methylthiazol-5-yl) ethyl phosphate (cTHz-P). HMP-PP and cTHz-P are then condensed into thiamine monophosphate (TMP) and finally phosphorylated to make the active cofactor TPP (15). Thiamine biosynthesis activities are found in bacteria, yeast, and plants. Transport proteins specific for thiamine, thiamine monophosphate (TMP), TPP, and other thiamine precursors are broadly distributed across all three domains of life (10, 19-23). Almost without exception, the genes and operons encoding these diverse thiamine transport and biosynthesis pathways are regulated by highly conserved TPP riboswitches.

Riboswitches are highly conserved *cis*-encoded and *cis*-acting regulatory RNAs found in the untranslated regions of messenger RNAs. Riboswitches bind small molecules, changing mRNA secondary structure and controlling expression outcomes at the level of transcription elongation, translation initiation, or RNA splicing (24, 25). TPP riboswitches are unique in that they are the only riboswitch found across all three domains of life (26). TPP riboswitches are so highly conserved, they are now used to identify thiamine biosynthesis and transport genes in diverse organisms (9). In bacteria, TPP riboswitches regulate biosynthesis and transport genes at either the transcriptional or translational level (27). TPP-bound forms of riboswitches that control transcription of downstream genes promote formation of a stem-loop that causes premature transcription termination. TPP riboswitches that act at the level of translation promote formation of a secondary structure that blocks ribosome binding when TPP is bound (28). TPP riboswitches that act at the level of transcription have been found primarily in Gram-positive bacteria, whereas Gram-negative bacteria tend to have TPP riboswitches that act at the level of translation (29). It has been noted that TPP riboswitches in *E. coli* that repress translation (when TPP is bound) also reduce mRNA stability, resulting in reduced steady-state levels of the transcript (30). While TPP riboswitch structures and functions have been extensively studied in the model organisms *E. coli* and *B. subtilis*, there has been little characterization of these conserved regulatory elements in other microbes, including those belonging to the phylum Bacteroidetes.

Computational analyses predicted that TPP riboswitches are present and regulate putative thiamine biosynthesis and transport pathways across the Bacteroidetes (9, 22). TPP riboswitch-controlled genes in *Bacteroides thetaiotaomicron*, a prominent member of the Bacteroidetes, have recently been characterized (10, 20). These studies have shown that both thiamine transport and biosynthesis are critical to the fitness of *B. thetaiotaomicron* under thiamine-limiting growth conditions. In this study, we build on previous work to investigate the global gene expression response of three representative *Bacteroides* species to exogenous thiamine and to determine the mechanisms of TPP riboswitch-dependent regulation of thiamine biosynthesis and transport genes. We found that each of the three species we studied had a distinct transcriptional response to exogenous thiamine. Investigation of the mechanisms of TPP riboswitch-mediated regulation of downstream genes and operons revealed that *B. thetaiotaomicron* uses both transcriptional- and translational-acting riboswitches, whereas *Bacteroides uniformis* and *Bacteroides vulgatus* use TPP riboswitches that act at the level of transcription. Identification of TPP riboswitches across the Bacteroidetes revealed a few different configurations of TPP riboswitch-controlled operons and suggested that both transcriptional and translational mechanistic classes are widely distributed.

## Results

### *Global transcriptional responses to exogenous thiamine vary between* Bacteroides *species*

Our previous study identified putative thiamine biosynthesis and transport genes in 114 *Bacteroides* genomes (10). We found biosynthesis and transport genes in 107 of 114 genomes, and a few genomes that appeared to contain either biosynthesis (2 of 114) or transport (5 of 114) genes. To further understand the diversity of thiamine biosynthesis and transport functions and regulation in *Bacteroides*, we chose three representative species to further characterize in this study. In *B. thetaiotaomicron*, we identified and characterized three gene clusters (Fig. 1A, 1C, 1E) involved in thiamine biosynthesis (*thiSEGCHF*_*Bt*_*-tenI*_*Bt*_), outer membrane thiamine transport (*OMthi*_*Bt*_) and cytoplasmic membrane transport (*pnuT*_*Bt*_*-tnr3*_*Bt*_) in a previous study (10). Each of these three gene clusters is preceded by a putative TPP riboswitch (Fig. 1A). In *B. uniformis*, we identified two putative TPP riboswitch-controlled gene clusters: for biosynthesis (*thiSEGCHF*_*Bun*_*-tenI*_*Bun*_) and transport (*OMthi*_*Bun*_*-pnuT*_*Bun*_*-tnr3*_*Bun*_) (Fig. 1C). *B. vulgatus* has only a single putative TPP riboswitch dependent operon for thiamine biosynthesis (*thiSEGCHF*_*Bv*_*-tenI*_*Bv*_) and apparently lacks thiamine transport genes (Fig. 1E).

**FIG 1.**
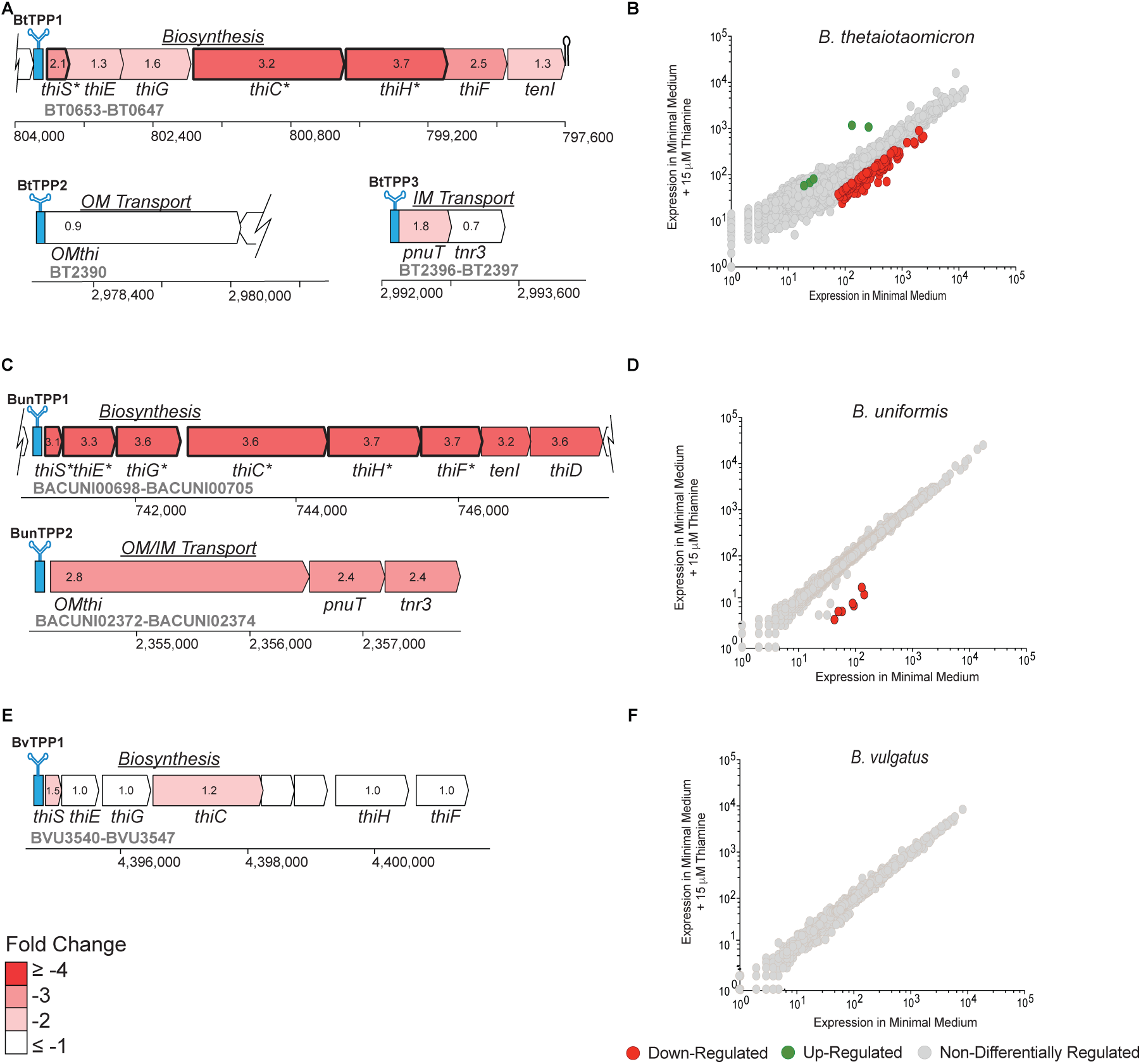
Transcriptional response to exogenous thiamine. Operons controlled by TPP riboswitches are shown for (A) *B. thetaiotaomicron*, (C) *B. uniformis*, and (E) *B. vulgatus*. Expression changes across operons are visualized in a schematic of each operon and color coded according to fold-change values comparing RNA-seq reads for cells grown with and without added thiamine. Negative fold-change values indicate that reads were reduced in cells grown in the presence of thiamine. Asterisks (*) and bold numbers indicate significance expression differences (*q-value* ≤ 0.05 and a change in expression ≥ 2-fold). All operons and distances are represented to horizontal scale. RNA-seq data from (B) *B. thetaiotaomicron* (previously in Costliow and Degnan 2017), (D) *B. uniformis*, and (F) *B. vulgatus* grown in a defined minimal medium without thiamine (0 nM thiamine added) and with thiamine (15 μM thiamine added). Grey circles indicate genes that are not differentially regulated. Green circles indicate genes that are significantly up regulated in the presence of thiamine. Red circles indicate genes that are significantly down regulated in the presence of exogenous thiamine. Data were analyzed using Rockhopper. The experiment was carried out in biological duplicate.

Our previous study showed that *B. thetaiotaomicron* differentially expresses genes comprising a number of pathways, including thiamine biosynthesis and transport, in the presence versus absence of exogenous thiamine (10). To determine if the transcriptional response to thiamine limitation is consistent across *Bacteroides* species we analyzed transcriptomes of *B. uniformis* and *B. vulgatus* in minimal medium with and without exogenous thiamine (as described in Materials and Methods and (10)). A global view of differential gene expression in the absence versus presence of exogenous thiamine is shown for *B. thetaiotaomicron* (Fig. 1B and (10)), *B. uniformis* (Fig. 1D) and *B. vulgatus* (Fig. 1F). Whole transcriptome responses to exogenous thiamine were markedly different among these three species. *B. thetaiotaomicron* had 151 genes significantly differentially expressed by ≥ 2-fold in the presence of exogenous thiamine compared to its absence (Fig. 1B). Genes that showed reduced expression in the presence vs. absence of exogenous thiamine in *B. thetaiotaomicron* encode functions belonging to a few pathways, including thiamine biosynthesis, amino acid biosynthesis, TCA cycle enzymes, purine, and pyruvate metabolism (Table S1 and 10). In contrast, minimal transcriptome responses to exogenous thiamine were observed in *B. uniformis* (Fig. 1D) and *B. vulgatus* (Fig. 1F). *B. uniformis* responded to exogenous thiamine with reduced expression of the genes in the TPP riboswitch-controlled thiamine biosynthesis cluster (*thiSEGCHF*_*Bun*_*-tenI*_*Bun*_, Fig. 1C, 1D). There were no significant differences in the gene expression profile of *B. vulgatus* in the absence vs. presence of exogenous thiamine (Fig. 1F). Our previous study revealed that *B. vulgatus* lacks an *OMthi* homolog found in many other *Bacteroides* species (10), suggesting that *B. vulgatus* may not be able to recognize or take up exogenous thiamine. In addition, while *B. vulgatus* does have a *pnuT* homolog, this gene is not preceded by a TPP riboswitch and is not clustered with other putative thiamine transport or biosynthesis genes, suggesting that this PnuT-like protein may have a different substrate specificity compared to the *B. thetaiotaomicron* and *B. uniformis* PnuT proteins (20, 31). Together, these data suggest that *B. vulgatus* does not respond transcriptionally to exogenous thiamine because it is not recognized or transported.

TPP riboswitches have been found to control either transcription elongation or translation initiation, depending on the organism (27). We noted that in *B. thetaiotaomicron* and *B. uniformis*, TPP riboswitch-controlled operons for biosynthesis and transport had different responses to exogenous thiamine at the level of mRNA abundance based on RNA-seq experiments. We hypothesized that this might reflect differences in TPP riboswitch-mediated mechanisms of regulation of these different operons. In *B. thetaiotaomicron*, only the biosynthesis operon mRNAs were differentially regulated in response to exogenous thiamine (Fig. 1A, (10)). The abundance of *OMthi*_*Bt*_ and *pnuT*_*Bt*_*-tnr3*_*Bt*_ mRNAs encoding transport functions in *B. thetaiotaomicron* was not significantly different in the presence or absence of exogenous thiamine (Fig. 1A, (10)), suggesting that the TPP riboswitches controlling *B. thetaiotaomicron* thiamine transport genes might act at the level of translation. The levels of TPP riboswitch-controlled biosynthesis and transport operon mRNAs in *B. uniformis* were reduced in the presence of exogenous thiamine (Fig. 1C). To confirm the results of RNA-seq with respect to thiamine-induced changes in mRNA levels, RT-qPCR was performed. Primers were designed to measure levels of biosynthesis (*thiC*), outer membrane TonB-dependent transporter (*OMthi*), and inner membrane transporter (*pnuT*) mRNAs in each organism. The mRNA abundance trends observed in all three organisms in RNA-seq were replicated in RT-qPCR experiments. In *B. thetaiotaomicron, thiC*_*Bt*_ mRNA levels were 60-fold higher in cells grown without thiamine compared to cells grown in the presence of 10 μM thiamine (Fig. 2A). The abundance of *OMthi*_*Bt*_ and *pnuT*_*Bt*_ mRNAs was equivalent in cells grown with and without thiamine. In *B. uniformis, thiC*_*Bun*_ mRNA levels were 2-fold higher in cells grown without thiamine compared to cells grown with thiamine (Fig. 2B). In contrast to *B. thetaiotaomicron, B. uniformis* transporter mRNA (*OMthi*_*Bun*_ *pnuT*_*Bun*_) levels also responded to exogenous thiamine with ∼3-fold higher levels in cells grown without thiamine (Fig. 2B). Transcript levels of the *B. vulgatus* TPP riboswitch-controlled biosynthesis operon did not respond to exogenous thiamine (Fig. 2C), also consistent with RNA-seq results. These data suggest that in *Bacteroides* organisms that have thiamine transport functions, exogenous thiamine affects transcript levels of thiamine biosynthesis genes. The transcriptional response to thiamine varied between species for mRNAs encoding transport functions.

**FIG 2.**
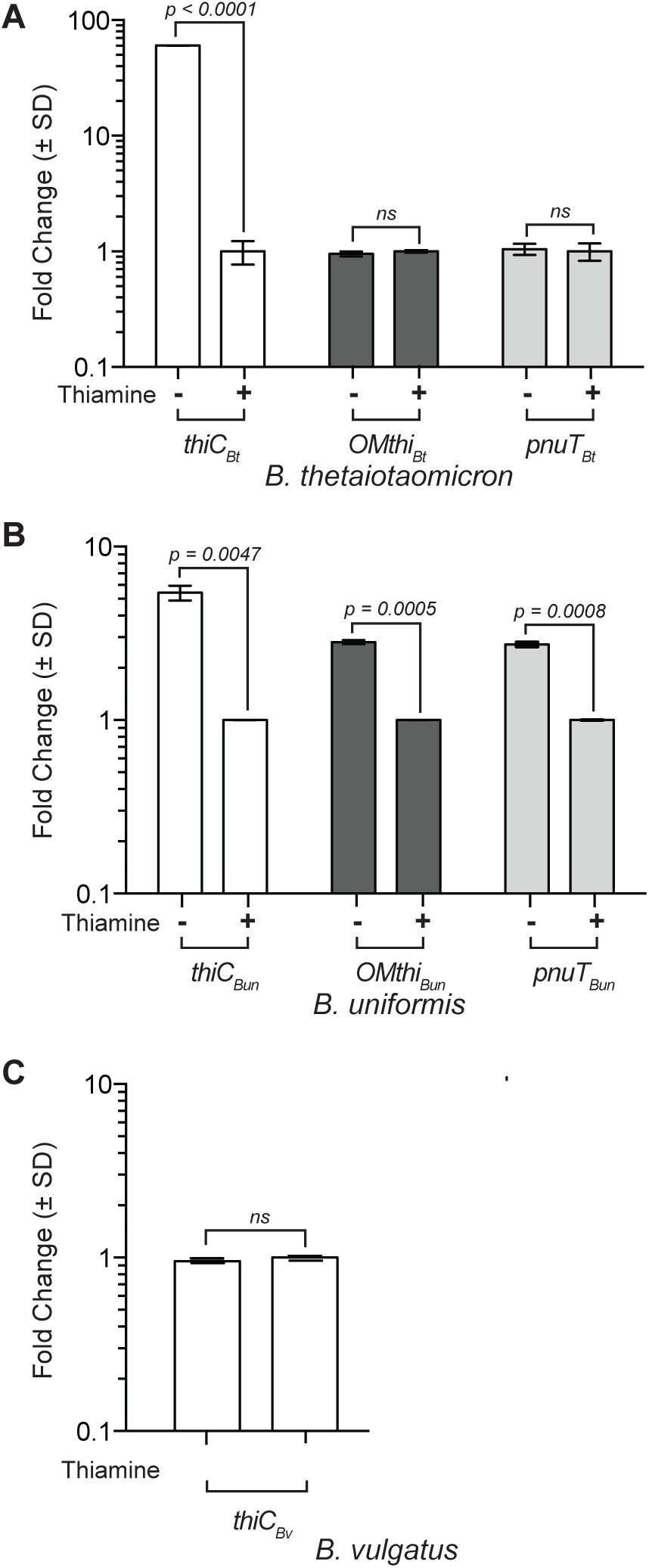
RT-qPCR confirmation of transcript level responses to exogenous thiamine. Transcript abundance of TPP riboswitch-regulated biosynthesis and transport genes was measured by RT-qPCR in (A) *B. thetaiotaomicron*, (B) *B. uniformis*, and (C) *B. vulgatus*. Primer sets were specific for genes indicated below bar graphs. Transcript levels of cells grown in the absence of thiamine (-) were normalized to levels in cells grown in the presence of 10,000 nM thiamine. Error bars represent the standard deviation (S.D.) for each experiment carried out in biological triplicate. Statistical significance for the differential abundance of gene transcripts were determined via a Student’s t-test. Changes in expression were considered significant when the fold change was ≥ 2-fold with a *p*-value ≤ 0.05.

### *Translational regulation of TPP-controlled operons in* Bacteroides

We next investigated if TPP riboswitch-controlled operons that did not show a transcriptional response to exogenous thiamine were instead regulated at the level of translation. We constructed nanoluciferase translational fusions to the start codon of the first gene in TPP riboswitch-controlled operons. All nanoluciferase (NanoLuc) fusions are controlled by native promoters and *cis*-acting TPP riboswitches. It is important to note that these translational fusions would capture thiamine-dependent transcriptional regulation caused by riboswitch-mediated premature transcription termination and translational regulation, *e*.*g*., riboswitch-mediated changes in ribosome binding site accessibility. Fusions for the three predicted TPP riboswitches (numbered as indicated in Fig. 1A) in *B. thetaiotaomicron* are called pBtTPP1, pBtTPP2 and pBtTPP3 (Fig. 3A). *B. uniformis* fusions were pBunTPP1 and pBunTPP2 (Fig. 3B), and the *B. vulgatus* fusion was called pBvTPP1 (Fig. 3C). Strains containing each fusion were grown in media containing a range of exogenous thiamine concentrations (from 0 to 10,000 nM). Promoterless NanoLuc controls were tested in each strain background without and with 10,000 nM thiamine. These controls showed that thiamine did not significantly alter luminescence (RLU/OD_630_), as indicated by dashed lines in Fig. 3A-D.

**FIG 3.**
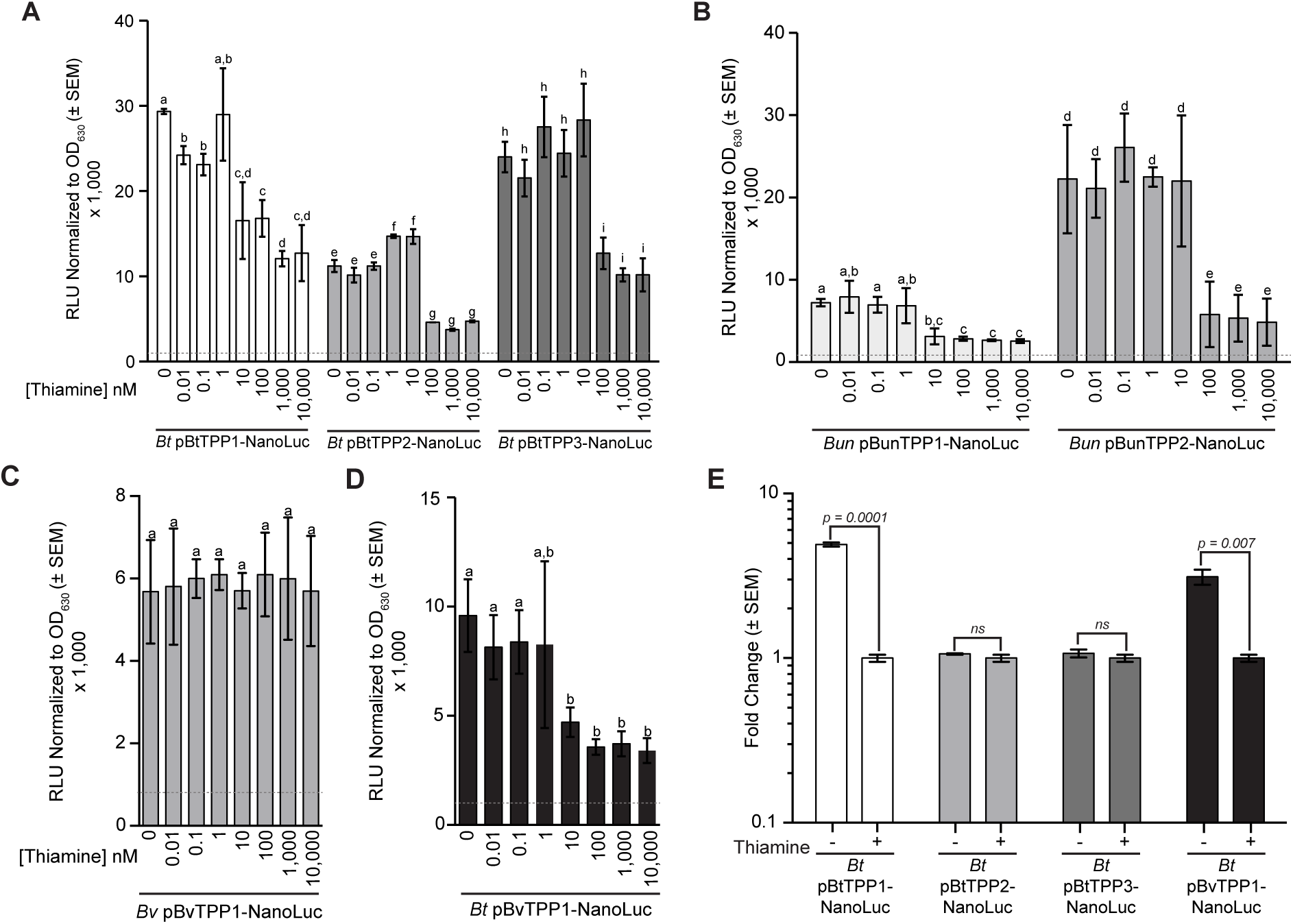
Nanoluciferase assays of *Bacteroides* TPP riboswitches. Nanoluciferase translational fusions to genes downstream of all six TPP riboswitches found in (A) *B. thetaiotaomicron*, (B) *B. uniformis*, and (C) *B. vulgatus* were tested. The *B. vulgatus* reporter fusion was integrated into the *B. thetaiotaomicron* chromosome (D) to test the mechanism of regulation in response to thiamine because *B. vulgatus* does not respond to exogenous thiamine. RT-qPCR was carried out to measure reporter fusion mRNAs in *B. thetaiotaomicron* with thiamine (+) or without thiamine (-) to confirm the mechanism of regulation by predicted riboswitches from *B. thetaiotaomicron* and *B. vulgatus* (E). Relative light units (RLU) were normalized to the OD_630_ of the bacterial cultures before cell lysis and activation of the nanoluciferase reaction. Cells were harvested in mid-log phase for all samples (OD_630_ between 0.4-0.6). The activity of each fusion was represented as the averaged RLU/ OD_630_ × 1000 from six independent biological replicates. Error bars represent the standard error of the mean (SEM). Letters (a, b, c, etc.) indicate significant differences based on a *p-value* ≤ 0.05 using a Student’s t-test. Grey dashed lines in A, B, C, and D indicate average RLU/ OD_630_ × 1000 from 3x biological replicates of a promoter-less nanoluciferase vector control for each experiment.

In *B. thetaiotaomicron*, the BtTPP1 reporter fusion was repressed as exogenous thiamine concentrations increased, with statistically significant differences beginning at 10 nM thiamine (Fig. 3A). This result, combined with the reduced levels of *thiSEGCHF*_*Bt*_*-tenI*_*Bt*_ mRNA observed in RNA-seq (Fig. 1B) and RT-qPCR (Fig. 2A), suggests that BtTPP1 riboswitch acts at the level of transcription. The *B. thetaiotaomicron* BtTPP2 and BtTPP3 riboswitches control thiamine transport *OMthi*_*Bt*_ and *pnuT*_*Bt*_*-tnr3*_*Bt*_ genes, and the levels of these mRNAs were not affected by exogenous thiamine (Figs. 1B and 2A). In contrast, the BtTPP2 and BtTPP3 reporters were repressed in the presence of exogenous thiamine, starting at 100 nM (Fig. 3A). These data suggest that the *B. thetaiotaomicron* riboswitches controlling transport operons regulate at the level of translation.

The reporters fused to *B. uniformis* riboswitches controlling biosynthesis (BunTPP1) and transport (BunTPP2) operons followed a pattern similar to that observed in *B. thetaiotaomicron*. The biosynthesis riboswitch reporter BunTPP1 was repressed by concentrations of exogenous thiamine at or above 10 nM, while the transport riboswitch reporter BunTPP2 required concentrations of 100 nM or greater for repression (Fig. 3B). These data, along with RNA-seq (Table S1) and RT-qPCR (Fig. 2B) are consistent with the model that both TPP riboswitches in *B. uniformis* act at the level of transcription.

In *B. vulgatus*, the riboswitch reporter controlling biosynthesis genes (BvTPP1) did not respond to exogenous thiamine (Fig. 3C). However, since it appears that *B. vulgatus* lacks a thiamine transport system, we reasoned that the lack of a response could be due to failure to take up and accumulate the thiamine provided exogenously. To test whether the BvTPP1 riboswitch could respond to thiamine in an organism competent for thiamine transport, we moved BvTPP1-NanoLuc into *B. thetaiotaomicron* (*Bt* pBvTPP1-NanoLuc, Fig. 3D). In a *B. thetaiotaomicron* background, the *B. vulgatus* BvTPP1 biosynthesis riboswitch was repressed at concentrations at or above 10 nM exogenous thiamine (Fig. 3D). This response is similar to both the *B. thetaiotaomicron* and *B. uniformis* thiamine biosynthesis regulating TPP riboswitches (Fig. 3A, B). The response to exogenous thiamine observed for the *B. vulgatus* TPP1 riboswitch reporter in *B. thetaiotaomicron* shows that the biosynthesis regulating riboswitch of *B. vulgatus* is functional. To determine whether the *B. vulgatus* TPP1 riboswitch controls transcription or translation of biosynthesis genes, we performed RT-qPCR on *B. thetaiotaomicron* strains carrying reporter fusions to detect changes in NanoLuc fusion mRNAs in the absence and presence of thiamine (Fig. 3E). As controls we also monitored levels of fusion mRNAs for the *B. thetaiotaomicron* transcriptional riboswitch (BtTPP1) and the translational riboswitches (BtTPP2 and BtTPP3). The results for controls were as expected – levels of the biosynthesis BtTPP1 reporter mRNA were reduced in the presence of thiamine, and levels of the transporter BtTPP2 and BtTPP3 fusion mRNAs were unchanged. The levels of *B. vulgatus* biosynthesis BvTPP1 reporter mRNA were also reduced in response to exogenous thiamine in the *B. thetaiotaomicron* strain background (Fig. 3E) suggesting that the *B. vulgatus* TPP riboswitch acts at the level of transcription.

Collectively, these data suggest *Bacteroides* species utilize riboswitches that act at either the transcriptional or the translational level. The riboswitches controlling biosynthesis operons are more sensitive than transport operon riboswitches, suggesting a hierarchical response to thiamine concentrations in the range of 10 to 100 nM.

### Distances of TPP Riboswitches Correspond to Regulatory Mechanism

Our results so far suggest that several TPP riboswitches – BtTPP1, BunTPP1, BunTPP2, and BvTPP1 – in *B. thetaiotaomicron, B. uniformis*, and *B. vulgatus* regulate downstream gene expression at the level of transcription. In contrast, BtTPP2 and BtTPP3 regulation was only observed at the level of translation. We observed that the transcriptionally acting riboswitches in *B. thetaiotaomicron, B. uniformis, and B. vulgatus* were located further upstream from the first gene in the operon compared to the translational-acting riboswitches in *B. thetaiotaomicron*. The differences in distance between the TPP riboswitch and the downstream gene that correlate with regulatory mechanism are consistent with previously characterized TPP riboswitches in model organisms (25, 29, 30). Transcriptional-acting riboswitches commonly found in Gram-positive bacteria are typically located >40-nt upstream of the start codon of the adjacent gene, whereas translational-acting riboswitches often found in Gram-negative bacteria are usually <30-nt from the downstream gene. These trends that are consistent with our observations in *Bacteroides* led us look more globally at the distance between TPP riboswitches and downstream genes across the Bacteroidetes.

We analyzed 219 TPP riboswitches in 114 Bacteroidetes genomes. The riboswitches were grouped and categorized according to the genes that they regulate (Fig 4A, Table 2). Among the 219 riboswitches identified, 122 were predicted to regulate thiamine biosynthetic operons. *Bacteroides* species have biosynthetic operons in which critical genes for thiazole synthesis, pyrimidine synthesis, and thiamine condensation are in the same TPP riboswitch-controlled operon (Fig. 4A). More distantly related Bacteroidetes, e.g., *Parabacteroides* and *Paraprevotella* species appear to have split biosynthesis operons, with one TPP-controlled gene cluster for thiazole synthesis and a second for HMP synthesis and TMP condensation (Fig. 4A). A total of 97 TPP riboswitches were found upstream of predicted thiamine transport genes, primarily operons with *OMthi* (85 of 97) or *pnuT* (12 of 97) as the riboswitch-proximal gene (Fig. 4A, Table 2). In contrast to the diversity observed among TPP riboswitch controlled thiamine biosynthesis operons; thiamine transport operon organization is highly conserved. TPP riboswitches regulate both the inner and outer membrane transporters in addition to a thiamine pyrophosphokinase (*tnr3*) (Fig. 4A).

**Table 1.**
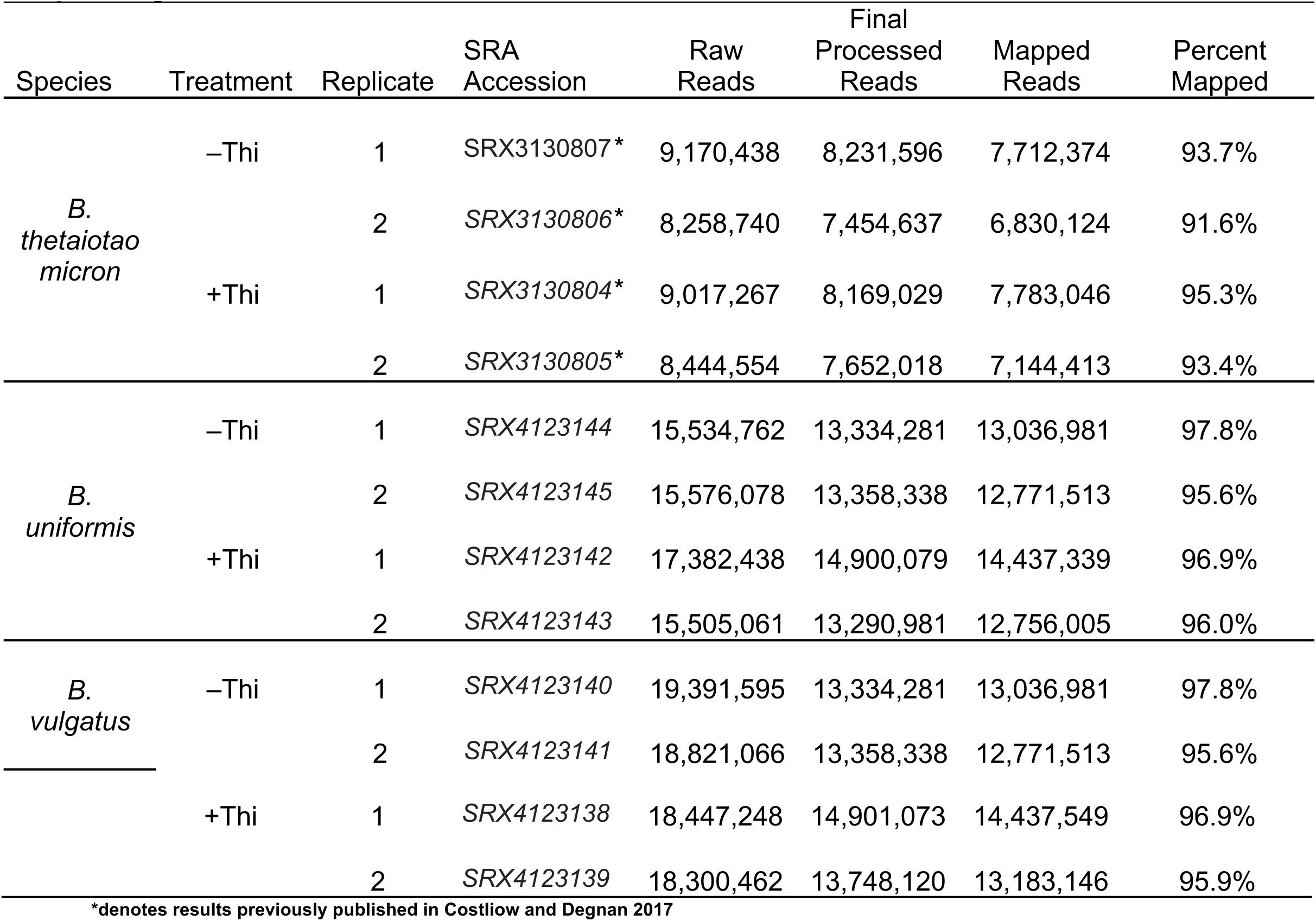
Reads of quality trimmed and mapped RNA sequencing.

**Table 2.**
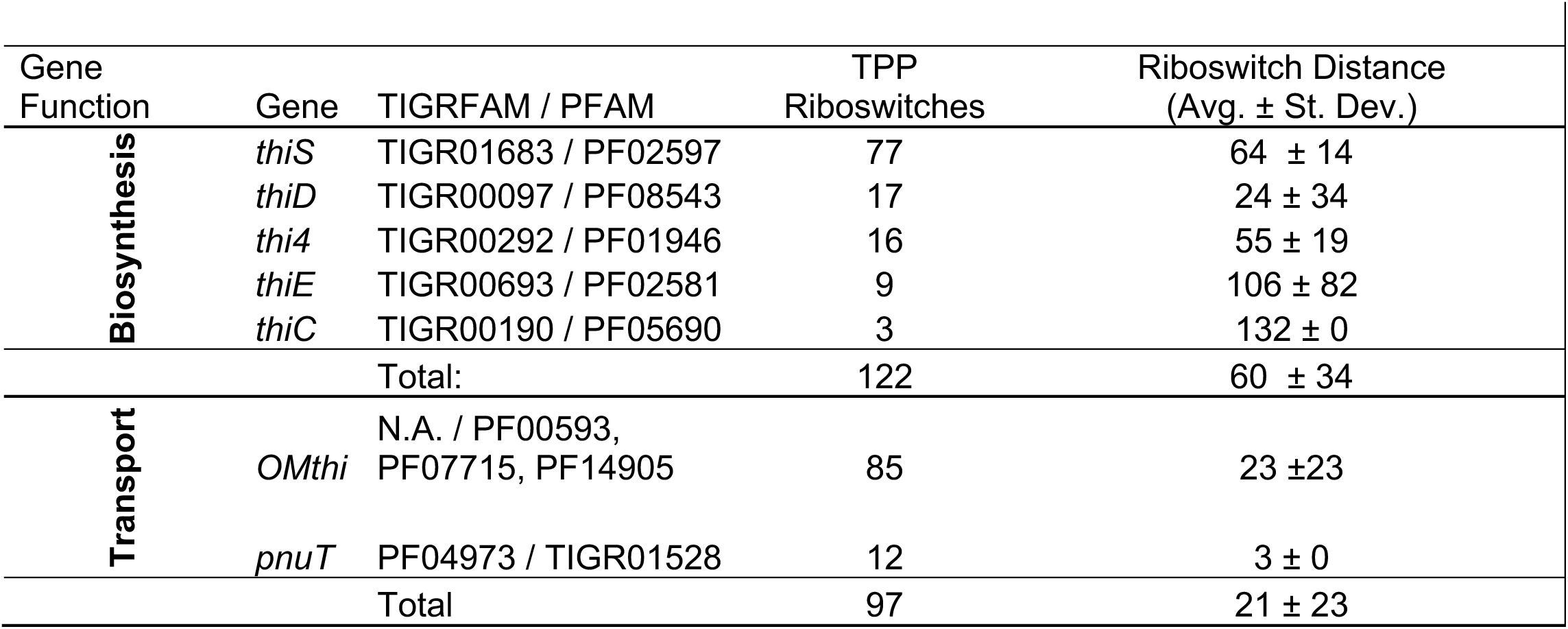
Average distance distributions of TPP riboswitches from the first gene of their regulon.

**FIG 4.**
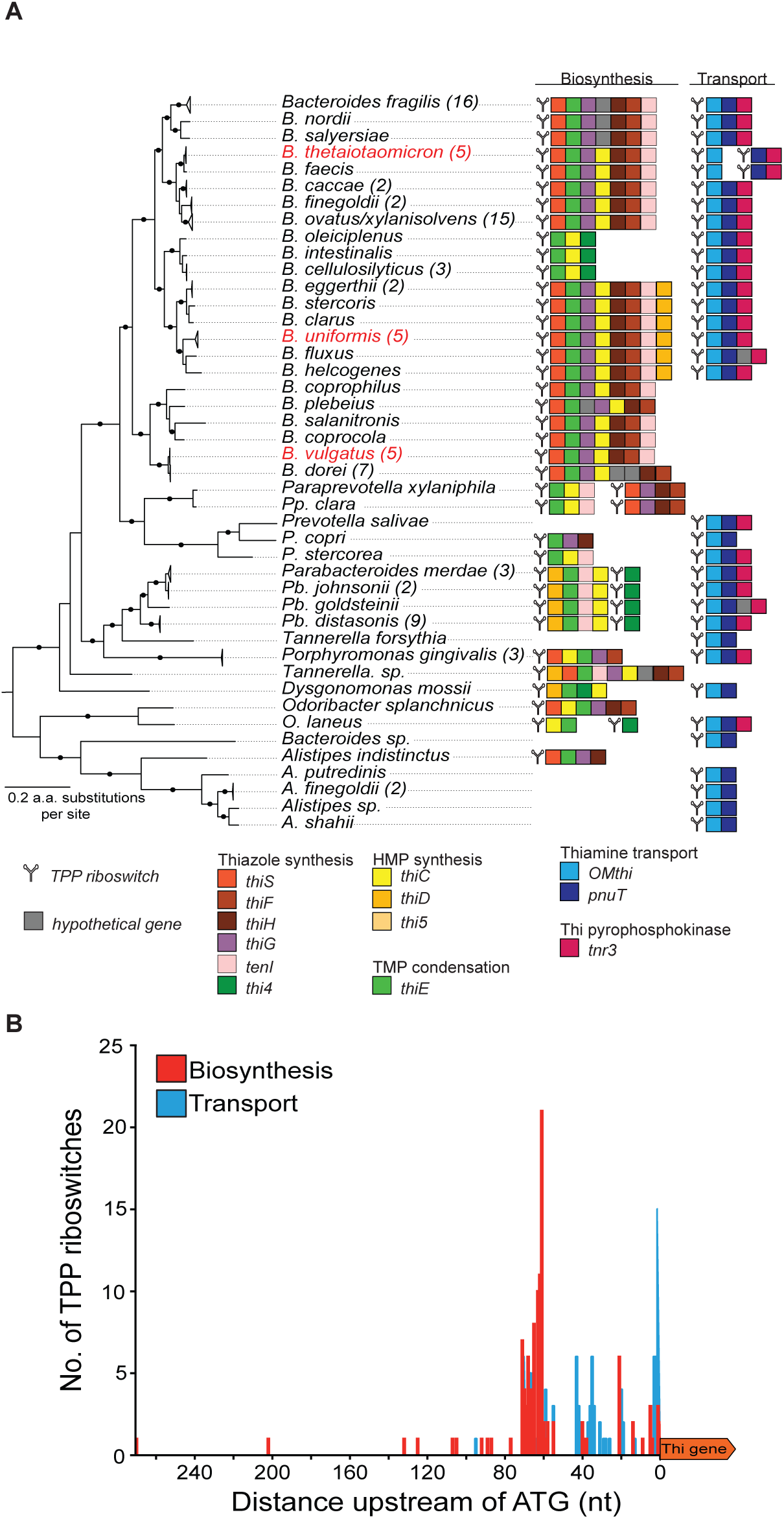
TPP riboswitches throughout the Bacteroidetes. Infernal and the RFAM 00059 covariance model were used to identify TPP riboswitches across the Bacteroidetes. (A) ORFs downstream were inspected to determine the operons directly preceded by putative TPP riboswitches. Function and syntenic context were assigned to all predicted riboswitch regulated operons. (B) The distance of each TPP riboswitch from its predicted operon was recorded and plotted and divided into the subcategories based on functional class of genes regulated by the riboswitch - thiamine biosynthesis (red) or thiamine transport (blue).

For the transcriptional-acting TPP riboswitches in *B. uniformis, B. vulgatus*, and *B. thetaiotaomicron* (biosynthetic operon only), we observed a >50-nt distance between the TPP riboswitch and the start codon of the first gene in the operon. In contrast, for the *B. thetaiotaomicron* thiamine transport operons, the TPP riboswitch was <5-nt from the first gene in each operon. We next analyzed the distances between TPP riboswitches and downstream genes more broadly across the Bacteroidetes (Table 2). While there is a range, the average distance between TPP riboswitches and downstream thiamine biosynthesis genes is 60 nt (± 34 nt). The average distance between TPP riboswitches and downstream thiamine transport genes is shorter at 21 nt (± 20 nt) (Table 2). We plotted the number of occurrences of riboswitches at specific distances from downstream genes, classified by the predicted function of the downstream operon (Fig. 4B). This analysis illustrated the overall pattern we observed, where TPP riboswitches upstream of biosynthesis genes were located further from the start codon and TPP riboswitches upstream of transporter genes were located closer to the start codon. Based on our experimental data from *B. thetaiotaomicron, B. uniformis*, and *B. vulgatus* and the observed TPP riboswitch distances across other gut Bacteroidetes, we predict that the TPP riboswitches located <20 nt from the start codon regulate at the level of translation while riboswitches >40 nt from the start codon regulate at the level of transcription.

## Discussion

Thiamine plays an essential role in both host and microbial metabolism, so thiamine biosynthesis and transport may play a key role in modulating the dynamics of host-microbe interactions. It is thought that most of the thiamine in the gastrointestinal tract is derived from the host’s diet (5, 23), and that microbial community composition may be influenced by competition for this key nutrient (10). In this study, we characterized variable TPP riboswitch-dependent regulation of thiamine biosynthesis and transport genes in three representative *Bacteroides* species. In all three organisms, the thiamine biosynthesis genes were controlled at the level of transcription by a TPP riboswitch located ∼50-nt upstream from *thiS*, the first gene in the biosynthesis operon. In *B. thetaiotaomicron*, the thiamine transport genes were controlled at the level of translation, by TPP riboswitches encoded immediately upstream of the start codons of transport genes. When we examined gut Bacteroidetes more broadly, we found that the distances between TPP riboswitches and downstream start codons were consistently correlated with the functional category, with longer distances for biosynthesis genes and shorter distances for transport genes. We postulate that in the Bacteroidetes, TPP riboswitches upstream of biosynthesis genes commonly control expression at the level of transcription elongation whereas riboswitches upstream of transport genes frequently control translation initiation.

Historically, it was thought that mechanisms of regulation by TPP-dependent riboswitches were divided between Gram-negative bacteria, where regulation of translation was common, and Gram-positive bacteria, where transcription attenuation was typical (29, 32, 33). Recent studies have demonstrated that these distinctions are not so clear. For example, the *lysC* riboswitch in *E. coli* is a dual-acting riboswitch. Ligand binding promotes both translational repression and RNase E-dependent cleavage and subsequent mRNA decay (30). Chauvier, *et* al., recently found a role for transcriptional pausing in modulation of TPP binding and subsequent regulatory events for the *E. coli thiC* riboswitch (34, 35). Pausing of the transcription complex at a regulatory site within the *thiC* riboswitch inhibits TPP binding and allows transcription elongation and translation. If TPP binds co-transcriptionally, translation is inhibited and Rho-dependent termination occurs. Our results further emphasize that diverse organisms use both transcriptional and translational strategies for regulation mediated by TPP-dependent riboswitches.

Trends in distances between TPP riboswitches and downstream genes in *B. thetaiotaomicron, B. uniformis* and *B. vulgatus* suggest that riboswitches located further upstream from the adjacent start codon control downstream gene expression at the level of transcription elongation. These more distant riboswitches were found upstream of thiamine biosynthesis genes in all three organisms and thiamine transport genes in *B. uniformis*, and all promoted a thiamine-dependent reduction in the respective mRNAs. It is possible that one or more of these riboswitches could act at both transcriptional and translational levels, or that translational regulation could be coupled to nuclease-dependent degradation. The riboswitches upstream of thiamine transport genes in *B. thetaiotaomicron* were directly adjacent to the downstream start codons, and these riboswitches clearly regulated translation in response to thiamine, but had no effect on transcript levels. The molecular details of TPP riboswitch-dependent regulation in *Bacteroides* species will require additional work. In these organisms, much remains to be discovered even in terms of basic mechanisms of regulation of transcription, translation, and mRNA processing and decay.

Our translational fusions revealed that TPP riboswitches regulating biosynthesis genes were responsive to thiamine at lower concentrations compared to riboswitches controlling transport genes. Biosynthesis genes were significantly downregulated in the presence of 10 nM thiamine, while transport genes responded only to concentrations at or above 100 nM. These concentrations are similar to the concentrations at which *E. coli* TPP riboswitches regulate cognate genes (33), suggesting that internal thiamine requirements may be similar across gut microbes. In addition, the 10 to 100 nM thiamine concentration range corresponds to the expected concentration of thiamine in the gut (20 to 2000 nM) (5). These data suggest that TPP riboswitch affinities among the Bacteroidetes and other gut bacteria may have evolved to operate at levels common to the human intestinal tract and to build a hierarchy of regulation between thiamine biosynthesis and transport to deal with rapidly changing nutrient availability in the gut. Such regulatory systems are not uncommon in the Bacteroidetes as a similar hierarchical regulatory phenomenon was observed for vitamin B_12_ riboswitches which respond to different concentrations of B_12_ to differentially control when transporters of varying affinities are expressed (36). It is worth noting that despite the similarities between thiamine and B_12_ riboswitches in hierarchical responses to concentrations of the cognate vitamin, the B_12_ riboswitches found in Bacteroidetes have only been observed to act at the level of translation (37).

We inspected the publicly available genomes to identify TPP riboswitches in 114 Bacteroidetes genomes and found that TPP riboswitches were commonly found upstream of putative thiamine biosynthetic and transport operons. These predicted riboswitches appear to follow the distance trends observed in *B. thetaiotaomicron, B. uniformis*, and *B. vulgatus*. If distances between the TPP riboswitch and downstream start codon are indicative of regulatory mechanism, as suggested by our results, then Bacteroidetes commonly use both transcriptional and translational acting TPP riboswitches to maintain internal thiamine homeostasis in the gut. While regulatory thresholds of Bacteroidetes TPP riboswitches may be similar across species and genera, global responses to thiamine are distinct. We have shown that *B. thetaiotaomicron* has a large suite of genes that are differentially regulated in the presence of exogenous thiamine, while closely related *B. vulgatus* and *B. uniformis* have no or a much narrower response. Such regulon differences have been observed in *E. coli* and *Salmonella enterica* where thiamine metabolism in the absence of thiamine show strikingly different global effects. In *E. coli*, branched-chain amino acid catabolism is involved while *S. enterica* leverages purine metabolism to produce precursors for thiamine synthesis (16-18).

Bacteroidetes TPP riboswitch structures are highly conserved across the species investigated we have investigated here, but we have shown that at least *B. thetaiotaomicron* utilizes both transcriptional and translational acting TPP riboswitches. While a single organism using different mechanisms for TPP riboswitch-dependent regulation is rare, it is not unheard of for a single riboswitch to transcriptionally regulate genes for biosynthesis of its ligand while regulating its transporter at the level of translation (25, 30). While the structure of the adaptor domain that binds TPP is highly conserved, the expression platform evolves much more quickly and likely has the largest impact on the mechanism of regulation mediated by each riboswitch (38). When riboswitches first evolved in the RNA world, it is believed that transcriptional control came first and allowed riboswitches to control the function of ribozymes (39). The regulation of translation likely evolved later and was retained because it is advantageous in some cases to have a reversible form of regulation to respond to rapidly changing environmental concentrations of the riboswitch ligand (38, 40). In the context of Bacteroidetes in the gut it may be beneficial to have transporters regulated at the level of translation as it is a reversible and fast-acting mechanism of regulation that could respond to rapidly changing thiamine concentrations in the gut. Conversely, the tighter control of thiamine biosynthesis in the Bacteroidetes via transcriptional regulation may help save the energy of producing multiple biosynthetic enzymes unless they are strictly necessary under very limiting thiamine concentrations. The existence of these regulatory hierarchies between transcriptional and translational acting riboswitches are uncommon in model organisms studied so far. Expansion of model systems to include a greater diversity of bacterial phyla may reveal new examples that regulate responses to vitamins, amino acids, or other nutrients involved in core metabolic pathways (24, 36, 27).

The study of how thiamine impacts microbial metabolism, global gene expression and microbial physiology may help us understand nutrition-dependent mechanisms responsible for modulating microbial community dynamics in the gastrointestinal tract. Our work demonstrates that *B. vulgatus* is effectively blind to exogenous thiamine, suggesting that it will not compete for available thiamine with the host or other gut microbes. Reliance on biosynthesis for production of essential vitamins may be a strategy used by certain species to establish a foothold in complex communities where availability of these molecules varies depending on diet and competition. The lack of thiamine transport capacity of *B. vulgatus* was not obvious based on existing genome sequence annotation of open reading frames, highlighting the importance of functional genomics and accurate annotation of cis-acting regulatory elements like riboswitches. Further characterization of mechanisms regulating thiamine acquisition and biosynthesis might lead to development of new strategies to modulate microbial community composition and metabolic function through targeted approaches controlling the activities of vitamin-dependent riboswitches (10, 41). These strategies could have a variety of exciting therapeutic applications that leverage the beneficial roles of microbial communities in human health and nutrition.

## Materials and Methods

### Bacterial Strains and Plasmids, Culturing and Genetic Manipulation

Bacterial strains and plasmids and oligonucleotides used for strain construction are described in Table S2.

Culturing of *Bacteroides* strains *B. thetaiotaomicron* VPI-5482, *B. uniformis* ATCC 8492, and *B. vulgatus* ATCC 8482 occurred anaerobically at 37°C in liquid tryptone yeast extract glucose (TYG) medium (42) or a modified minimal medium with 0 to 10 μM (final concentration) of thiamine hydrochloride as described previously (10, 43, 44) or Difco brain heart infusion (BHI) agar with the addition of 10% defibrinated horse blood (QuadFive, Ryegate, MT, USA). Cultures were grown in a vinyl anaerobic chamber with an input gas mix consisting of 70% nitrogen, 20% carbon dioxide, and 10% hydrogen (Coy Laboratory Products, Grass Lake, MI, USA). *E. coli* S17-1 λ *pir* strains used for cloning and conjugation of reporter vectors were grown in LB medium at 37°C aerobically. Antibiotics were added to the media when appropriate at final concentrations as follows: ampicillin 100 μg/mL, gentamicin 200 μg/mL, and erythromycin 25 μg/mL.

Nanoluciferase reporter vectors were constructed via traditional cloning of the nanoluciferase gene (45, 46) into the vector pNBU2 (36, 47). DNA fragments encompassing TPP riboswitches and the first 5 codons of the downstream gene were amplified from genomic DNA by PCR using HiFi Taq MasterMix (KAPA Biosystems, Wilmington, MA, USA) and inserted in frame preceding the nanoluciferase gene in the pNBU2 vector. The constructs were transformed into conjugation donor *E. coli* S17-1 *λ pir*, and confirmed via PCR amplicon size and sequencing. After confirmation, vectors were then conjugated into *B. thetaiotaomicron* VPI-5482, *B. uniformis* ATCC 8492, and *B. vulgatus* ATCC 8482 making a single insertion into the genome of each strain (36, 47). A complete list of primers and plasmids used in this study are provided in Table 3.

### Expression analysis

Replicate cultures of wild-type *B. uniformis* and *B. vulgatus* were grown overnight in 5 mL minimal medium supplemented with a final concentration of 10 μM thiamine HCl (THI, > 99% pure) (Sigma Aldrich, St. Louis, MO, USA). Aliquots of each culture were pelleted by centrifugation (1 min at 13,300 x g), culture supernatants were decanted and the cells were washed 4 times in minimal medium with 0 nM thiamine. Cells were subcultured at a dilution of 1:2000 into 10 mL of minimal medium with either 0 or 15 μM (final concentration) thiamine HCl in two biological replicates. Cell growth was monitored and cells were harvested between OD_600_ 0.4 and 0.6 on a UV spectrophotometer. Total RNA was extracted using the Qiagen RNeasy kit and stored at −80°C. Total RNA was DNAse treated with DNase I (Thermo Fisher, Waltham, MA, USA) and NEB DNase treatment protocol. DNAse treated RNAs were re-purified using the Qiagen RNeasy kit (Hilden, Germany), quantitated using a Qubit 2.0 (Life Technologies, Carlsbad, CA, USA), and stored at −80°C. This was the same method used for a previously published RNA-seq experiment for *B. thetaiotaomicron* (10). RNA was submitted for integrity analysis, ribosomal RNA depletion, and library construction at the W.M. Keck Center for Comparative and Functional Genomics at the University of Illinois at Urbana-Champaign. RNA sequencing was performed using an Illumina 2500 HiSeq generating 100 nucleotide paired end reads for all samples.

For targeted gene expression analysis, total RNA was purified as described above and used to generate cDNA libraries using the first strand cDNA synthesis kit (Thermo Fisher, Waltham, MA, USA). RT-qPCR was performed on a Biorad CFX connect instrument (Biorad, Hercules, CA, USA) and SYBR Fast MasterMix 2x Universal (Kapa Biosystems, Wilmington, MA, USA) following the manufacturer’s instructions for triplicate biological samples with three technical replicates. Five probe sets were used in three strains to amplify products specific to: 16S ribosomal RNA, *thiC, thiS, OMthi*, and *pnuT*. 16S rRNA primers were used as the control (36) and novel primers for *thiC, thiS, OMthi*, and *pnuT* for *B. thetaiotaomicron* (10), *B. uniformis*, and *B. vulgatus* were designed using Primer3 (48). Standard curves were used to evaluate the efficiency of the amplification, all 5 genes had R^2^ ≥ 0.98 and slope between −3.30 and −3.40. Relative expression changes were calculated using the ΔΔC_q_ method (49).

### Nanoluciferase Reporter Fusion Assays

Reporter assays were performed through a modified and combined protocol based on Lim *et al*. 2017 and Mimee *et al*. 2015. *Bacteroides* strains with integrated pNBU2 TPP riboswitch nanoluciferase reporters were grown in a modified minimal medium and then washed and back diluted to an OD_600_ of 0.004 in 1.5 mL Axygen deep 96-well plates (Corning, Inc., Corning, NY) over a gradient of thiamine HCl (Sigma-Aldrich, St. Louis, MO). Growth was monitored by via a BioTek Synergy HTX Multi-Mode microplate reader (BioTek, Winooski, VT) (10). When cells reached mid-log phase growth (0.35-0.58 OD_600_) growth was stopped. 1 mL of mid log phase culture was spun down in the Axygen deep well plate at 3000 x g for 10 minutes and medium was poured off the pellets. Cell pellets were then resuspended in 50 μL of 10X BugBuster protein extraction reagent (MilliporeSigma, Darmstadt, Germany). Cell suspensions were then incubated at room temperature with gentle shaking for 10 minutes. Nanoluciferase extracts were mixed 1:1 (10 μL:10 μL) with Nano-Glo Luciferase mixture. Mixtures were then incubated for 5 minutes at room temperature. Luminescence was read on a BioTek Synergy HTX Multi-Mode microplate reader (BioTek, Winooski, VT). Relative Luminescence Units were normalized to the OD_600_ of cell cultures. Comparisons were made across thiamine concentrations using pairwise Student’s *t* tests in GraphPad Prism v6, and *P* values of ≤0.05 were considered significant.

### In Silico *identification of TPP riboswitches in* Bacteroides *species*

Detection of TPP riboswitches among Bacteroidetes was done as described in (10). Infernal 2.0 was used to identify riboswitches using the covariance model of TPP riboswitches in RFAM (RF00059) (50, 51) in 114 Bacteroidetes genomes available in RefSeq from the Human Microbiome Project (52, 53). To confirm that Infernal correctly predicted TPP riboswitches, TIGRFAM, PFAM, and Hidden Markov Models with HMMR were used to identify thiamine pyrophosphate biosynthesis and transport genes among the same 114 RefSeq Bacteroidetes genomes (53-55).

To analyze the trend in distances between riboswitches and downstream genes, nucleotide distances between coordinates identified with Infernal 2.0 and TIGRFAM, PFAM, and HMMR were taken and subtracted from one another (51, 54-57). Annotations and genes were then manually inspected utilizing Artemis to confirm that genes and riboswitches were called correctly and that distances were accurate and agreed with previous predictions.

## ACCESSION NUMBER

RNAseq data generated for this project have been submitted to the NCBI SRA database as study SRP148918

## ACKNOWLEDGEMENTS

We thank members of the Degnan and Vanderpool labs at the University of Illinois at Urbana-Champaign for comments on the manuscript. In addition, we thank Dr. Andrew Goodman and Dr. Bentley Lim from Yale University for protocols and plasmids associated with the NanoLuciferase reporter system. We also thank Alvaro Hernandez from the W.M. Keck Center for Comparative and Functional and Genomics at the University of Illinois at Urbana-Champaign for their technical services.

